# The Mitochondrial Ubiquitin Ligase MARCHF5 Cooperates with MCL1 to Inhibit Apoptosis in KSHV-Transformed Primary Effusion Lymphoma Cell Lines

**DOI:** 10.1101/2024.09.23.614413

**Authors:** Prasanth Viswanathan, Justine R. Bersonda, Jackson Gill, Alyssandra Navarro, Allison C. Farrar, Daniel Dunham, Karl W. Boehme, Mark Manzano

## Abstract

Kaposi’s sarcoma-associated herpesvirus (KSHV) causes several malignancies in people with HIV including Kaposi’s sarcoma and primary effusion lymphoma (PEL). We have previously shown that PEL cell lines require myeloid cell leukemia-1 (MCL1) to inhibit apoptosis. MCL1 is an oncogene that is amplified in cancers and causes resistance to chemotherapy regimens. MCL1 is thus an attractive target for drug development. The emerging clinical relevance and therapeutic potential of MCL1 motivated us to study the roles of this oncogene in PEL in depth. Using a systems biology approach, we uncovered an unexpected genetic interaction between MCL1 and MARCHF5 indicating that they function in the same pathway. MARCHF5 is an E3 ubiquitin ligase most known for regulating mitochondrial homeostasis and antiviral signaling, but not apoptosis. We thus investigated how MCL1 and MARCHF5 cooperate to promote PEL cell survival. CRISPR knockout (KO) of MARCHF5 in PEL cell lines resulted in a significant increase in apoptosis despite the presence of MCL1. The anti-apoptotic function of MARCHF5 was dependent on its E3 ligase and dimerization activities. Loss of MARCHF5 or inhibition of the 26S proteasome furthermore stabilized the MCL1 antagonist NOXA without affecting levels of MCL1. Interestingly, NOXA KO provides a fitness advantage to PEL cells suggesting that NOXA is the pro-apoptotic signal that necessitates the anti-apoptotic activities of MCL1 and MARCHF5. Finally, endogenous reciprocal co-immunoprecipitation experiments show that MARCHF5 and NOXA are found in the same protein complex. Our findings thus provide the mechanistic link that underlies the genetic interaction between MCL1 and MARCHF5. We propose that MARCHF5 induces the degradation of the MCL1 antagonist NOXA thus reinforcing the pro-survival role of MCL1 in these tumor cells. This newly appreciated interaction of the MCL1 and MARCHF5 oncogenes may be useful to improve the design of combination therapies for KSHV malignancies.

## INTRODUCTION

Kaposi’s sarcoma-associated herpesvirus (KSHV) is a human gammaherpesvirus that establishes life-long latency primarily in B cells. While the majority of infections remain asymptomatic, KSHV-positive people with HIV (PWH) can develop several hematologic diseases, including the lymphoproliferative disorder multicentric Castleman disease, and the rare but aggressive primary effusion B cell lymphoma (PEL) [1, 2]. PEL classically presents as effusions in body cavities, including the pleural, pericardial, and peritoneal spaces, although extra-cavitary cases have been reported [3, 4]. PEL comprises 4-8% of non-Hodgkin lymphoma cases in PWH [5, 6]. Currently, there is no standard therapy for PEL. Patients receive similar chemotherapy regimens as with aggressive non-Hodgkin lymphomas (cyclophosphamide, hydroxydauomycin/doxorubicin, Oncovine/vincristine, and prednisolone or CHOP) and anti- retroviral therapy (ART) [1]. Nevertheless, the median overall survival time is 6-22 months [3, 5–10].

While mutations from primary tumors have been reported, the majority of cases do not have genetic alterations in the tumor suppressor genes *TP53* or *RB1,* nor do they have *MYC* translocations frequently observed in other lymphomas [2–4, 11–14]. Instead, a major driver of PEL is oncogenic “addiction” or constitutive dependency on host genes that are transcriptionally or post-transcriptionally dysregulated by KSHV genes. In our previous genome-wide CRISPR- Cas9 knockout (KO) screens, we identified 210 PEL-specific oncogenic dependencies (PSODs) or addictions that are preferentially required by PEL cell lines but not by non-PEL cancer cell lines [15]. These PSODs function in viral persistence strategies that overlap with hallmarks of cancer: metabolic rewiring, transcriptional reprogramming, cell cycle progression, sustaining proliferative signaling, immune evasion, and inhibition of cell death. Because of their essential roles in tumor cell survival, PSODs are attractive therapeutic candidates to treat PEL.

At the top of this list was the anti-apoptotic protein myeloid cell leukemia-1 (MCL1), which ranked as the second most essential PSOD [15, 16]. *MCL1* is an oncogene that has copy number amplification in solid tumors and hematologic malignancies, including NHL [17]. Overexpression of MCL1 in different cancers is associated with disease severity and poor prognosis, underscoring the pro-survival role of this oncoprotein [18, 19]. With the development of the first-in-class MCL1 inhibitor S63845 [20], new and more potent MCL1 inhibitors are now being evaluated in the clinic [21]. This growing attention to MCL1 as a chemotherapeutic target to treat different cancers initiated our interest in the role of this oncogene in PEL.

MCL1 is a member of the B cell lymphoma-2 (BCL2) protein family that functions to prevent intrinsic apoptosis. BCL2 proteins are anchored to the mitochondrial outer membrane (MOM) to sequester monomers of the cell death effectors BAX or BAK1 [22]. The binding of BCL2 proteins to BAX/BAK1 prevents the monomers from assembling into a pore complex and initiating the apoptosis cascade. During intrinsic apoptosis, pro-apoptotic BCL2 homology region 3 (BH3)-only proteins are upregulated and bind with high affinity to the BCL2 proteins. This derepresses the inhibition of BAX/BAK1 and triggers MOM permeabilization (MOMP) [23]. While some BH3-only proteins can bind to all BCL2 family members (e.g., BIM, truncated BID, and PUMA), others are more selective. For instance, NOXA specifically neutralizes MCL1, while HRK has high specificity to BCL-XL.

We previously showed that PEL primary tumors and cell lines highly express MCL1 [15, 24]. PEL cell lines are only addicted to MCL1 and do not depend on the other pro-survival BCL2 genes [15]. Consistent with this finding, we and others demonstrated that inhibitors against MCL1, but not other BCL2 proteins, are highly effective in killing PEL cell lines [24–26].

Moreover, the KSHV latency-associated nuclear antigen (LANA) sequesters the MCL1 E3 ubiquitin ligase FWB7, thereby stabilizing the MCL1 protein and promoting tumor cell survival [26]. Together, these observations underscore the role of the oncogenic addiction to MCL1 in this viral lymphoma.

Given the importance of MCL1 in PEL, we sought to uncover mechanistic insights as to how MCL1 is regulated. In this study, we used an unbiased systems biology approach to identify that another high-scoring PSOD MARCHF5 cooperates with MCL1 function. We show that this mitochondrial E3 ubiquitin ligase targets the MCL1 antagonist NOXA for degradation and consequently reinforces MCL1 function in protecting tumor cells from apoptosis.

## RESULTS

### MCL1 and MARCHF5 are co-essential oncogene addictions in PEL cells

To gain unbiased insights into functions of MCL1, we took a systems biology approach to identify the genetic interactions of MCL1. We reanalyzed publicly available data from the Cancer Dependency Map (DepMap) [27]. DepMap integrates data from CRISPR screens from >1000 human cancer cell lines that identify genes essential for cell fitness to create “co-essentiality” maps. Genes that function in the same pathway have positive interactions because their individual knockout have similar phenotypic effects (co-essential genes or synergists). On the other hand, genes that have opposing functions have antagonistic interactions because the knockout of one negates the phenotype caused by the knockout of the other (antagonists).

Because co-essentiality mapping only measures phenotypes and is thus agnostic to prior knowledge of gene function, this approach can lead to the discovery of new gene functions and pathway relationships [27–30].

We examined the co-essentiality maps of *MCL1* from DepMap. Consistent with its known role in preventing intrinsic apoptosis, *MCL1* shows antagonistic interactions with several pro-apoptotic effectors, including *PMAIP1* (encodes for NOXA), *BAX*, *APAF1*, and *BAK1* (**Fig. 1A, Table S1**). Similarly, *MCL1* exhibits co-essentiality with other members of the BCL2 family, *BCL2L2* and *BCL2* (ranks 3 and 11, respectively). However, the top co-essential gene with *MCL1* was the mitochondrial E3 ubiquitin ligase *MARCHF5*, which had a correlation score that is twice as strong as compared to *BCL2L2 or BCL2*. In addition, the E2 ubiquitin ligase *UBE2J2* ranked second, just before the BCL2 genes. Conversely, *MCL1* was the top co-essential gene of *MARCHF5,* followed by *UBE2J2* (**Fig. 1B, Table S1**). This relationship is surprising because MARCHF5 functions as a regulator of mitochondrial homeostasis and antiviral signaling but not apoptosis [31–33]. Together, these findings underscore the functional link between MCL1 and the mitochondrial ubiquitin ligase complex.

**Figure 1.**
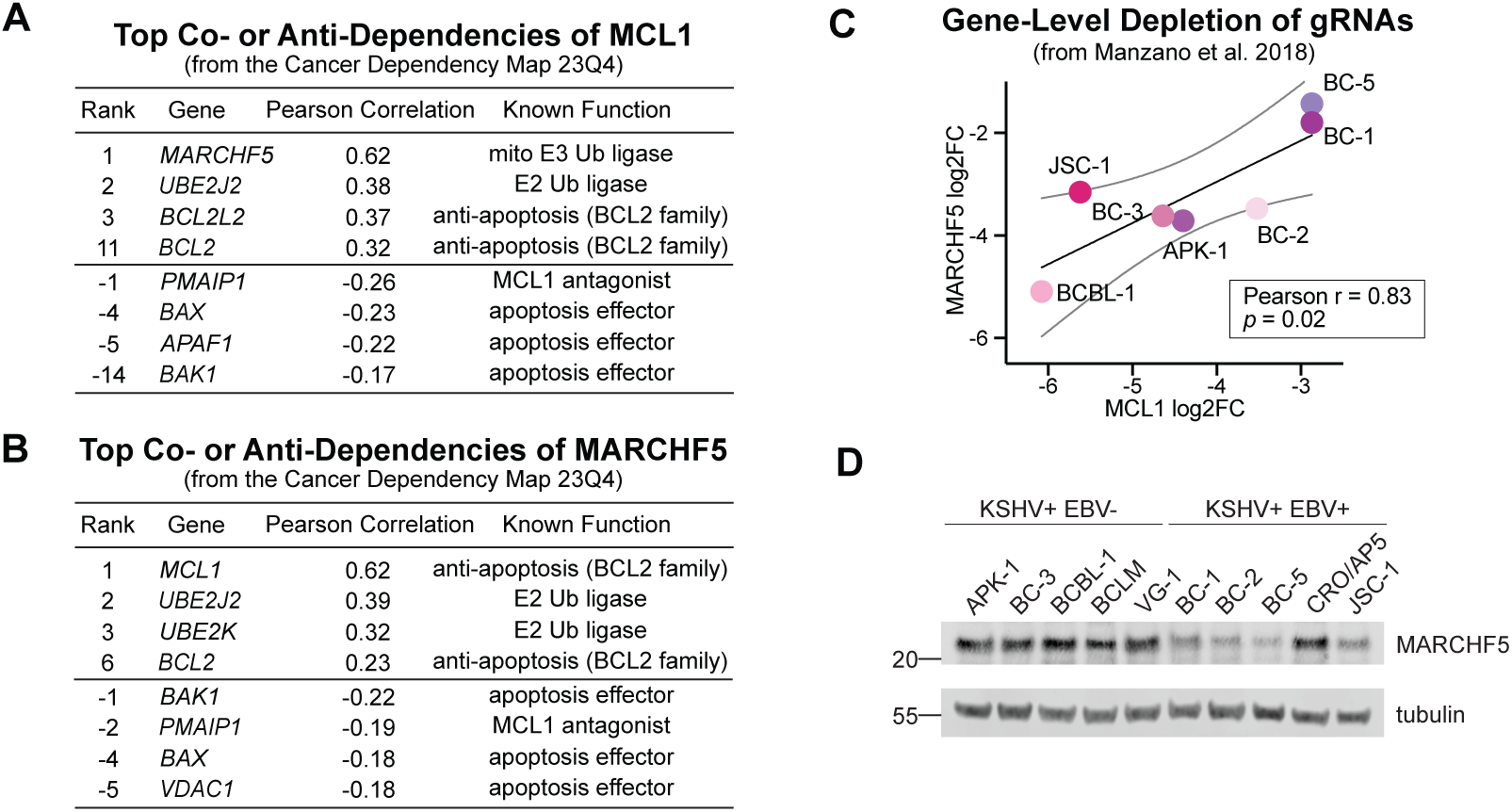
***A-B***. Top co- and anti-dependencies of ***A***. *MCL1* or ***B***. *MARCHF5* from the Cancer Dependency Map 23Q4. The top 100 genes are listed in Table S1. ***C***. Median fold changes of gRNAs targeting *MCL1* or *MARCHF5* in our gene essentiality CRISPR screens in PEL cell lines [15]. ***D***. Western blots of MARCHF5 in a panel of PEL cell lines that are infected with KSHV only or co-infected by KSHV and EBV.

To investigate if *MCL1* and *MARCHF5* exhibit co-essentiality in PEL cell lines, we reexamined our previous CRISPR screens that identify PSODs [15, 16]. In these screens, *MCL1* and *MARCHF5* ranked as the second and third strongest PSOD for PEL cell lines. Consolidated depletion scores for guide RNAs (gRNAs) targeting *MCL1* or *MARCHF5* strongly correlated across the individual cell lines, even more consistent than the computed correlation coefficient from DepMap (Pearson *r* = 0.83, **Fig. 1C**). In addition, MARCHF5 protein was expressed at similar levels in a wider set of cell lines (**Fig. 1D**). Together, co-essentiality analyses of CRISPR screens suggests that the two mitochondrial PSODs *MCL1* and *MARCHF5* may cooperate to promote fitness and survival of PEL cell lines.

### MARCHF5 is required for the survival of PEL cell lines

To validate if PEL cell lines are addicted to *MARCHF5* expression, we targeted *MARCHF5* for CRISPR-Cas9 KO using two independent gRNAs in two representative PEL cell lines: BC-3 expressing a constitutive Cas9 [15] and BCBL-1 expressing a doxycycline (Dox)-inducible Cas9 (iCas9) [34]. In both cell lines, MARCHF5 KO resulted in lower numbers of total live cells compared to the negative gRNA, which targets the safe harbor locus *AAVS1* (**Fig. 2A-B**).

**Figure 2.**
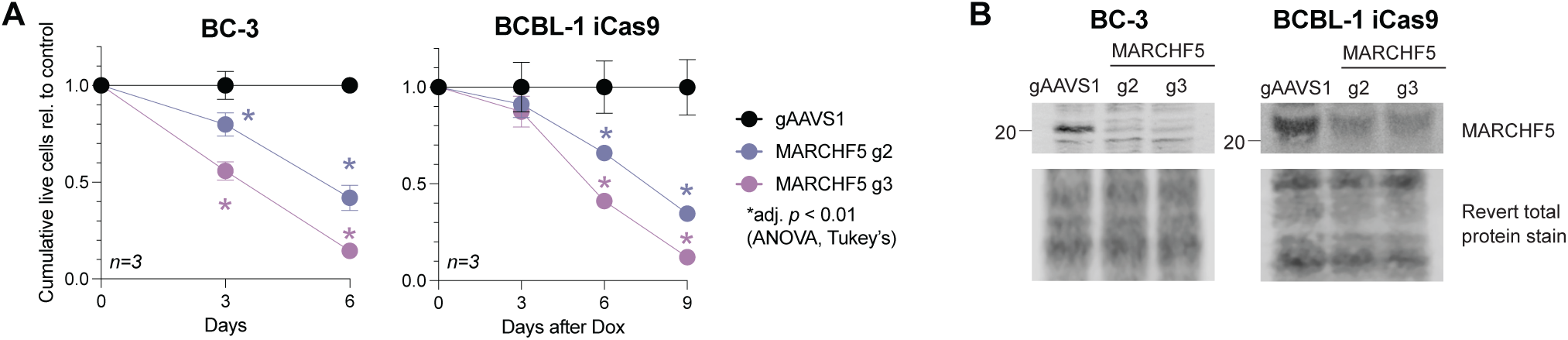
***A.*** Cumulative live cell counts (relative to control gAAVS1) of BC-3 Cas9 or BCBL-1 iCas9 cells expressing gRNAs targeting *MARCHF5* or *AAVS1*. ***B***. Western blots of MARCHF5 from cells in panel ***A***. Statistical differences were calculated using two-way ANOVA with Tukey’s multiple comparison post-hoc test (n=3). Adjusted *p* values reflect post-hoc comparisons between gAAVS1 to either MARCHF5 gRNA at the specific time point. Error bars, standard error of the mean.

The co-essentiality of *MARCHF5* and *MCL1* suggests that these mitochondrial proteins are functioning in the same pathway. To test whether MARCHF5 also prevents intrinsic apoptosis like MCL1, we measured the activities of the terminal caspases 3/7 using a luminescence-based Caspase-Glo 3/7 Assay. In both BC-3 Cas9 and BCBL-1 iCas9 cells, *MARCHF5* KO significantly increases caspase 3/7 activities (**Fig. 3A**). Similarly, levels of the early apoptotic marker Annexin V increased in *MARCHF5* KO cells (**Fig. 3B**). Taken together, these data confirm that MARCHF5 is dependency factor in PEL cell lines required to inhibit apoptosis.

**Figure 3.**
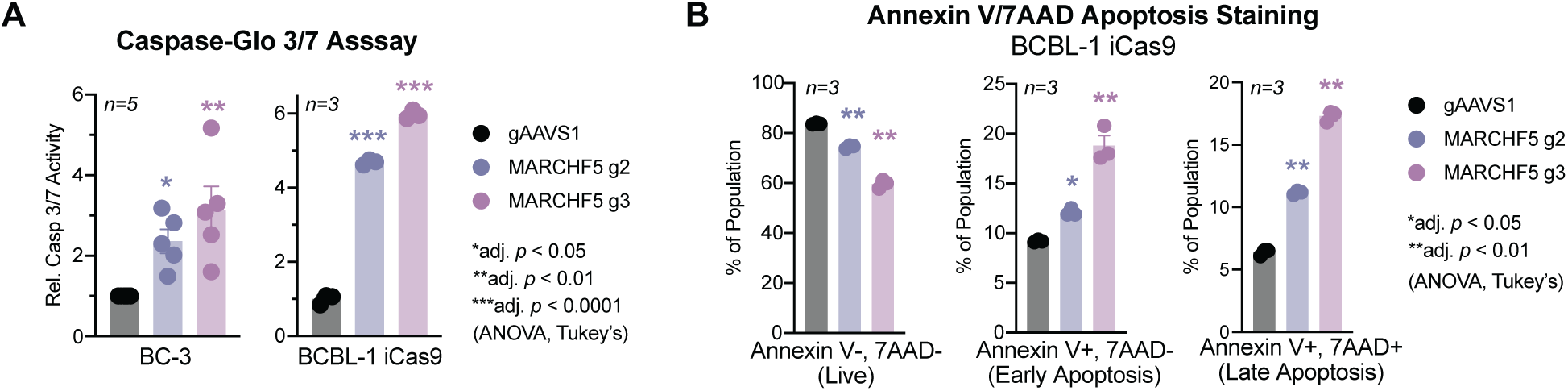
***A***. Western blots of MARCHF5 or FLAG in BCBL-1 iCas9 cells expressing gRNAs targeting *MARCHF5* or *AAVS1* and MARCHF5 cDNAs. ***B***. Cumulative live cell counts (relative to control gAAVS1) of BCBL-1 iCas9 cells in panel ***A*** after the addition of Dox. Statistical differences were calculated using two-way ANOVA with Tukey’s multiple comparison post-hoc test (n=6-8). Adjusted *p* values reflect post-hoc comparisons between gAAVS1 to either MARCHF5 gRNA at the specific time point. Error bars, standard error of the mean.

### Ubiquitin ligase activity and dimerization of MARCHF5 are required for the survival of PEL cell lines

MARCHF5 is an E3 ubiquitin ligase anchored to the mitochondrial outer membrane [31, 32]. It is known to regulate mitochondrial structure and dynamics by ubiquitinating proteins that regulate either mitochondrial fission or fusion. Furthermore, MARCHF5 protein levels are tightly regulated by auto-ubiquitination and homodimerization through GxxxG motifs to maintain proper mitochondrial homeostasis [35].

To determine if dimerization and/or the ubiquitin ligase activity is essential for MARCHF5 function in PEL cells, we transduced BCBL-1 iCas9 cells expressing the *MARCHF5* gRNAs with lentiviruses expressing cDNAs encoding for mCherry (negative control), a positive control 3XFLAG-tagged codon-optimized wild-type MARCHF5 (coMARCHF5 WT), a ligase mutant (coMARCHF5 H43W), or a dimerization mutant (coMARCHF5 4GS) [35, 36]. Codon optimization disrupts targeting sites for the *MARCHF5* gRNAs, thus restricting editing to only the endogenous alleles (**Fig. 4A**).

**Figure 4.**
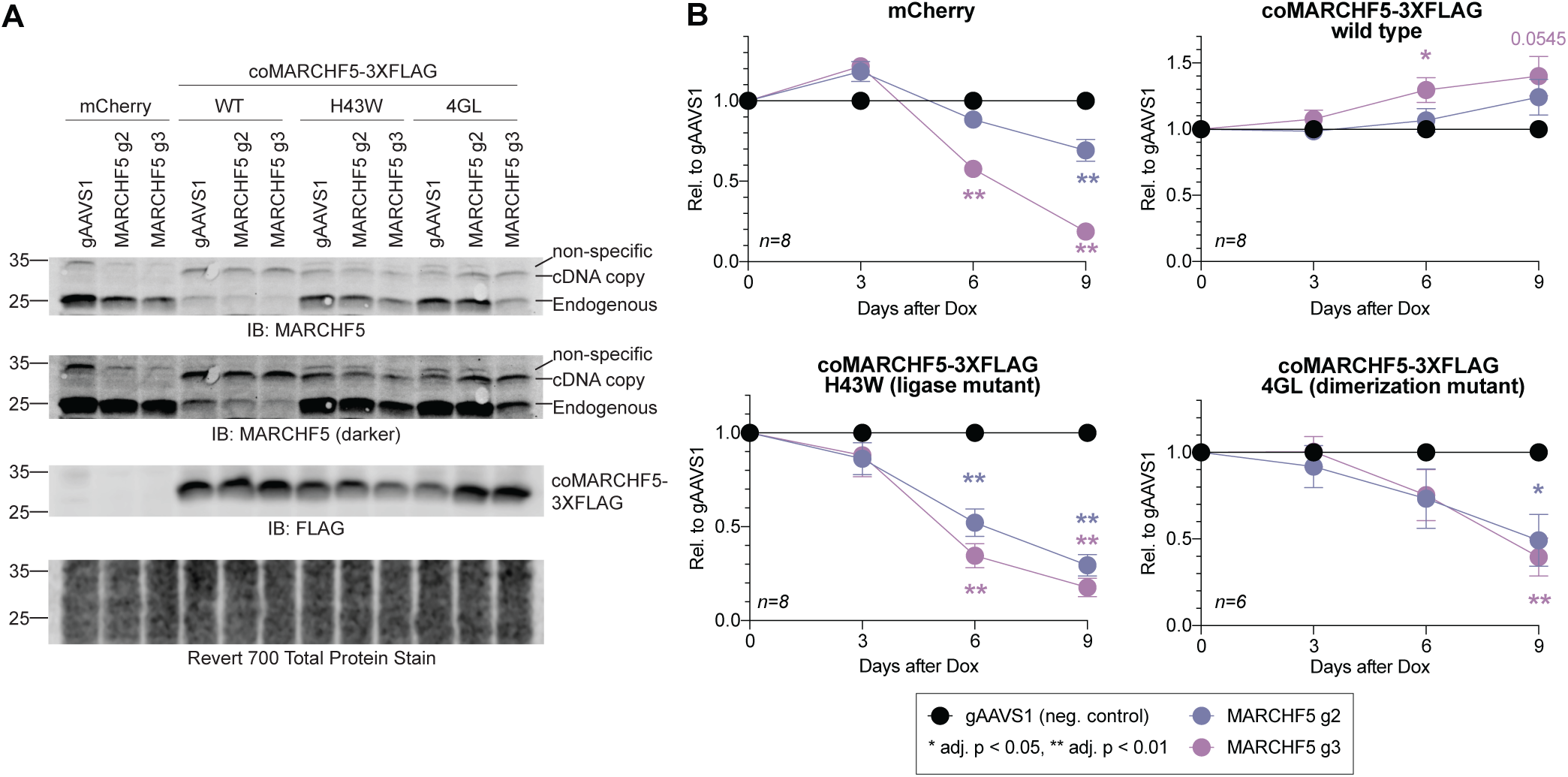
***A.*** Apoptotic activity measured by Caspase-Glo 3/7 Assay or ***B***. Annexin V-7AAD staining of cells from Fig. 3. Statistical differences were calculated using one-way ANOVA with Tukey’s multiple comparison post-hoc test (n=3-5). Adjusted *p* values reflect post-hoc comparisons between gAAVS1 to either MARCHF5 gRNA. Error bars, standard error of the mean.

Although the expression of the cDNA constructs was under the control of a strong CMV enhancer-promoter, these lentiviral copies were expressed to a lower level than the endogenous MARCHF5 (**Fig. 4A**). We noted that expression of coMARCHF5 WT also reduced endogenous protein. We previously observed these reductions in protein expression when attempting to over-express cDNAs in PEL cells. When endogenous proteins have low to moderate basal levels, we see high amounts of exogenous proteins [24]. However, when the endogenous protein is already expressed at high levels, the introduction of a transgene results in either modest expression of the transgene [24] or overexpression of the transgene but downregulation of the endogenous copy [37, 38]. This finding suggests that PEL cells have an auto-regulatory mechanism that restricts the expression of a protein to a maximum, whether it is expressed from an endogenous or exogenous locus or whether it contains an epitope tag.

Nevertheless, even though coMARCHF5 WT is only modestly expressed and reduced endogenous protein levels (**Fig. 4A**), the total MARCHF5 protein was sufficient to completely rescue the decrease of total live cells in *MARCHF5* g2 or g3 cells (**Fig. 4B**). This result indicates that the defects on cell fitness are specific to *MARCHF5* KO and not from off-target effects.

Moreover, neither the H43W ligase mutant nor the 4GL dimerization mutant rescued *MARCHF5* KO cells. These data collectively show that both the ubiquitin ligase activity and dimerization of MARCHF5 are required for the survival of PEL cell lines.

### Cell death mediated by *MARCHF5* KO is not caused by MAVS activation or lytic replication

One of the described functions of MARCHF5 is to negatively modulate antiviral signaling [33]. Upon sensing viral infections, RIG-I-like receptors (RLRs) are activated and bind to the signaling adapter MAVS on the mitochondrial surface. MAVS consequently transduces RLR signaling by activating antiviral kinases. This eventually leads to the phosphorylation of the transcription factors IRF3 and IRF7 and the transcription of interferon-stimulated genes. MAVS-mediated signaling is kept in check through a negative feedback loop wherein MARCHF5 induces MAVS protein turnover. Because MAVS restricts KSHV lytic replication [39], it is possible that the cell death from loss of MARCHF5 results from altered MAVS homeostasis and, consequently, the latency-lytic balance.

To test if cell death from *MARCHF5* KO is caused by dysregulated MAVS levels and signaling, we immunoblotted for total MAVS, phosphorylated IRF3 (p-IRF3), and total IRF3. Total MAVS, p-IRF3, and IRF3 remained unchanged in BC-3 *MARCHF5* KO cells (**Fig. 5**). Moreover, we did not detect upregulation of representative early and lytic proteins (K-bZIP and ORF26, respectively). Similar results were observed in BCBL-1 iCas9 cells. These data indicate that *MARCHF5* KO-induced apoptosis is not due to MAVS stabilization or aberrant lytic reactivation.

**Figure 5.**
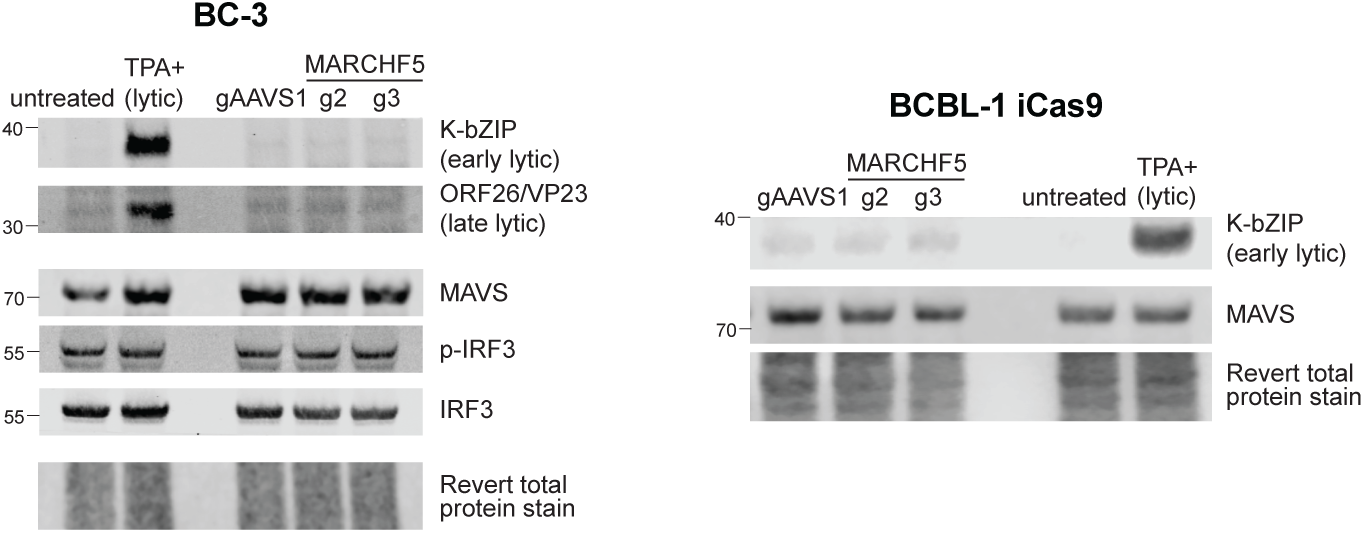
Western blots of lytic proteins and MAVS signaling pathway in wildtype or *MARCHF5* KO BC-3 Cas9 or BCBL-1 iCas9 cells.

### NOXA restricts the steady-state survival of PEL cell lines

Thus far, our current and previous work [15, 24] has shown that PEL cells exhibit oncogenic co-dependencies to MARCHF5 and MCL1 to prevent intrinsic apoptosis, independent of inhibition of MAVS signaling or lytic reactivation. However, it remains unclear how they are mechanistically connected. We hypothesize that MARCHF5 prevents apoptosis by ubiquitinating an antagonist of MCL1 for proteasome degradation such as the BH3-only proteins. During apoptosis, BH3-only proteins neutralize BCL2 family proteins to initiate the oligomerization of BAX/BAK1. The BH3-only proteins exhibit different affinities to the BCL2 family with some being more promiscuous binders (e.g. BIM, PUMA, BAD, BMF) while others (NOXA and HRK) are highly selective (**Fig. 6A**). Co-dependency interactions from DepMap reveal that both *MCL1* and *MARCHF5* are strongly anti-correlated with *PMAIP1* (NOXA) which supports our hypothesis that MARCHF5 ubiquitinates an MCL1 antagonist for degradation (**Figs. 1A-B, Table S1**).

**Figure 6.**
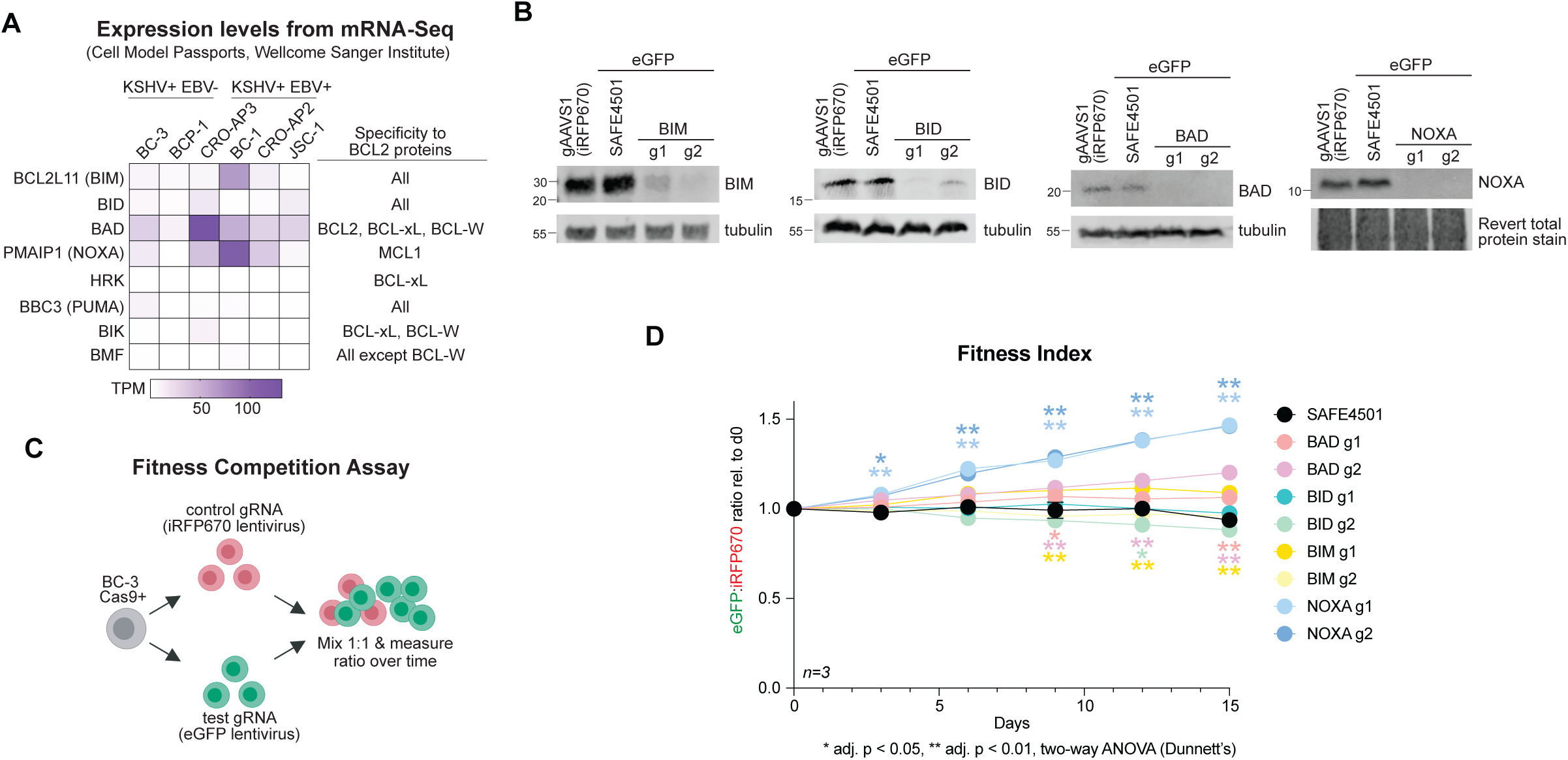
***A***. mRNA expression of BH3-only proteins in PEL cell lines from Cell Model Passports. ***B***. Western blots of BH3-only proteins that were targeted for Cas9 KO in BC-3 cells. ***C***. Experimental flow chart of the Fitness Competition Assay. BC-3 Cas9 cells were transduced with a lentivirus encoding the control gAAVS1 and an iRFP670 fluorescent protein or a second lentivirus encoding the targeting gRNA and an eGFP reporter. Equal amounts of control iRFP670 cells were mixed with targeted eGFP cells at the beginning of the growth curve. Fitness indices as reflected by the ratio of eGFP to iRFP670 (eGFP:iRFP670) were measured over 15 days by flow cytometry. ***D***. Fitness indices for each BH3-only KO competition relative to day 0. Statistical differences were calculated using two-way ANOVA with Dunnett’s multiple comparison post-hoc test (n=3). Adjusted *p* values reflect post-hoc comparisons between gSAFE4501 to BH3-only protein gRNA. Error bars, standard error of the mean.

To identify the relevant BH3-only protein that is constitutively transducing the apoptotic signal in PEL cells that necessitates the anti-apoptotic activities of MCL1 and MARCHF5, we measured the relative fitness of BC-3 cells with wildtype or Cas9-targeted BH3-only proteins. We hypothesized that Cas9 KO of the apoptotic trigger would provide a fitness advantage to these tumor cells. Because there are at least eight BH3-only proteins, we first narrowed our candidate list to the top four genes that had measurable basal RNA expression in different PEL cell lines: *BCL2L11* (BIM), *BID*, *BAD*, and *PMAIP1* (NOXA) (**Fig. 6A**). We then targeted each of these genes for KO in BC-3 Cas9 cells using two independent test gRNAs expressed from a lentivirus encoding an eGFP fluorescence reporter (**Fig. 6B**). As a negative control, we used the SAFE4501 gRNA that targets a non-functional, non-genic region [40]. In parallel, we derived another negative control cell line expressing the gAAVS1 gRNA from a lentivirus encoding an iRFP670 fluorescence reporter. We then performed a Fitness Competition Assay wherein eGFP+ cells expressing the test gRNA were mixed at equal ratio (1:1) with the negative control iRFP670+ cells expressing gAAVS1 (**Fig. 6C**). Relative fitness of the KO cells was monitored using the ratio of eGFP to iRFP670 as a readout by flow cytometry. Of the eight test KO cell lines, only NOXA KO cells had a measurable fitness advantage for both gRNAs over the negative control cells gSAFE4501 and gAAVS1 (**Fig. 6D**). These experiments suggest that NOXA is the major constitutive apoptotic signal that restricts the steady state survival of PEL cell lines.

### MARCHF5 is found in the same complex as the MCL1 antagonist NOXA

To test if NOXA is constitutively regulated at the protein level to promote the survival of PEL cells, we treated PEL cell lines with the proteasome inhibitor MG132. MG132 treatment dramatically stabilized NOXA while having marginal to no effects on MCL1 and MARCHF5 (**Fig. 7A**). This indicates that the NOXA undergoes rapid turnover by the proteasome at steady-state. To demonstrate that MARCHF5 facilitates NOXA turnover, we immunoblotted for NOXA in *MARCHF5* KO cells. *MARCHF5* KO cells had higher NOXA similar to MG132 treatment (**Fig. 7B**) suggesting that this E3 ligase is involved in the proteasomal turnover of NOXA. Finally, to determine if MARCHF5 directly interacts with NOXA, we performed co-immunoprecipitations (co-IP). IP of endogenous MARCHF5 complexes in BC-3 cells pulled down NOXA while reciprocal co-IP of NOXA complexes conversely pulled down MARCHF5 (**Fig. 7C**). These biochemical experiments confirm that MARCHF5 and NOXA form complexes in PEL cells. In sum, our data support the model that MARCHF5 cooperates with MCL1 by promoting the degradation of the MCL1 antagonist NOXA.

**Figure 7.**
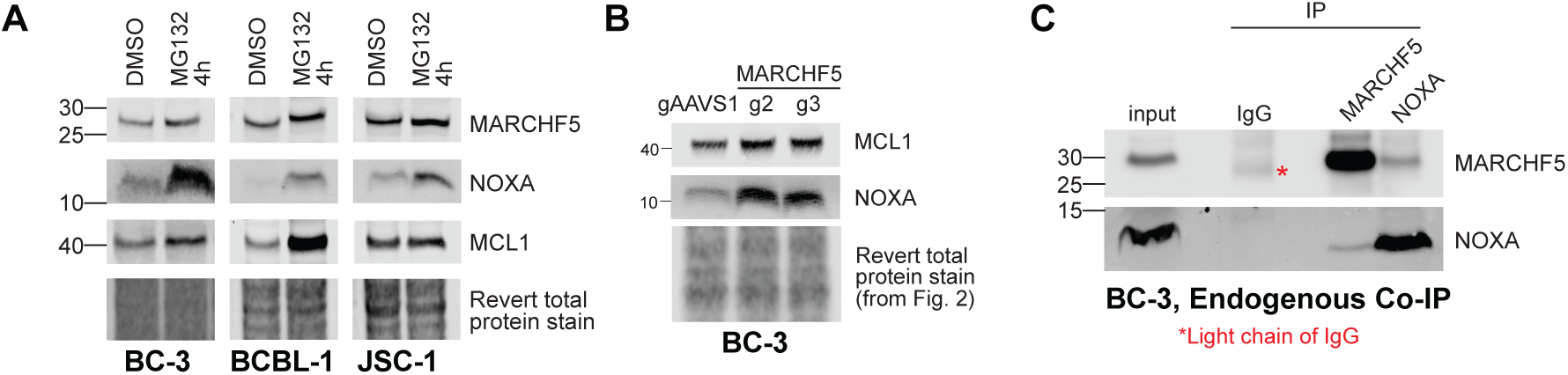
***A***. Western blots of PEL cell lines were treated with 1.25 μM MG132 for 4 hours. ***B***. Western blot for MCL1 and NOXA in wildtype or *MARCHF5* KO BC-3 Cas9. These protein lysates are the same ones used in Fig. 2 and thus use the same loading control. ***C***. Reciprocal co-immunoprecipitation (co-IP) of the endogenous MARCHF5-NOXA complex in BC-3 cells.

## DISCUSSION

In this study, we sought to uncover the genetic interactions of the anti-apoptotic protein *MCL1*, a clinically relevant oncogene in multiple cancers and KSHV-driven PEL [15, 17–21, 24]. Co-essentiality mapping from DepMap and our own CRISPR screens in PEL cell lines unexpectedly identified that MCL1 exhibits the strongest genetic interaction with the mitochondrial-resident E3 ubiquitin ligase MARCHF5, despite MARCHF5 not being known for being a canonical regulator of apoptosis. Strikingly, MCL1 and MARCHF5 are ranked back-to-back as the second and third strongest PSODs in PEL cell lines underscoring this co-dependency. We find that the amount of MCL1 protein is insufficient to protect from cell death because single CRISPR KO of MARCHF5 also results in apoptosis. Mechanistically, we find that MARCHF5 induces the turnover of NOXA protein thereby reinforcing the pro-survival role of MCL1 in these tumor cells.

The dependence on both MCL1 and MARCHF5 oncogenes underscores that PEL cell lines constantly require NOXA to be kept in check to protect from cell death. More recently, MARCHF5 has been implicated in the NOXA-mediated turnover of MCL1 itself [41]. MARCHF5 KO stabilizes both MCL1 and NOXA proteins indicating that MARCHF5 is targeting the entire MCL1-NOXA complex. However, we do not see this in PEL cells. MARCHF5 KO results in pronounced stabilization of NOXA but not MCL1. Moreover, MG132 treatment of different cell lines does not lead to MCL1 protein stabilization except for BCBL-1 cells suggesting that MCL1 is not being significantly regulated by proteasomal turnover. Our data thus point to a different mechanism by which MARCHF5 regulates NOXA separately from the MCL1-NOXA complex.

It remains unclear what causes the high levels of NOXA at steady-state that necessitates the addictions to both MCL1 and NOXA. It is worth noting that most PEL cell lines have functional p53 and low but detectable activated p53 [13, 42]. It is possible that latent infection and/or replicative stress induced by proliferating tumor cells causes this low-level activation of p53.

Thus, p53-responsive genes like NOXA need to be controlled by other mechanisms such as MARCHF5-mediated degradation of NOXA and the LANA-mediated stabilization of MCL1. Interestingly, KSHV encodes for a lytic viral homolog of the BCL2 family (vBCL2) [43, 44].

Biochemical studies demonstrate that vBCL2 most resembles MCL1 in sequence and function [45]. Thus, it appears that a major strategy of KSHV to protect infected cells from cell death is to reinforce MCL1 activity or to bring in an MCL1-like protein.

Our study does not rule out that MARCHF5 may have other essential functions in PEL cells. It is possible that the addiction to MARCHF5 also involves the regulation of mitochondrial dynamics to meet the altered energy demands of these actively proliferating tumor cells. Moreover, MARCHF5 has recently been linked to mediating both peroxisome biogenesis and turnover [46, 47]. Given that peroxisome proteins are induced by the latency genes and are required for survival by KSHV-infected endothelial cells [48], it is also possible that the dependency to MARCHF5 facilitates peroxisomal metabolism. However, these functions are difficult to observe in PEL due to the confounding effects of apoptosis in subcellular structures. Future studies on the role of MARCHF5 in mitochondrial or peroxisome homeostasis in PEL cell lines should therefore be done in the absence of the apoptotic machinery such as using *BAX/BAK1* double knockout cells.

Finally, the co-essential relationship of MCL1 and MARCHF5 may be exploited for combination therapy to treat PEL. While MCL1 inhibitors or proteasome inhibitors effectively kill PEL cell lines *in vitro* and *in vivo* [15, 24, 26, 49], their use as monotherapies has limitations in the clinic. Two recent Phase I trials using MCL1 inhibitors show that these treatments result in elevated cardiac troponin and cardiovascular risk in patients [50, 51]. In addition, although bortezomib is being used to treat multiple myeloma patients, it also induces lytic reactivation in PEL cell lines [49, 52] which may pose complications in PWH. By targeting the MCL1-MARCHF5 axis using combination therapy with MCL1 inhibitors and bortezomib, it is possible that a synergistic effect can be achieved at lower doses. More effective treatments will likely prevent the development of resistance and induce apoptosis faster before the virus can complete the lytic cycle. In addition, the lower doses may prevent dose-dependent cardiotoxicity associated with MCL1 inhibitors.

Thus, factoring co-essentiality interactions of oncogenes in rational therapy design for KSHV malignancies may improve therapeutic strategies for these cancers.

## MATERIALS AND METHODS

### Cells and treatments

All cloning was performed in *E. coli* Stbl3 cells (Invitrogen, Carlsbad, CA). PEL cell lines were maintained in RPMI-1640 Medium with L-glutamine and sodium bicarbonate (Millipore-Sigma, St. Louis) supplemented with 10% or 20% Serum Plus II Medium Supplement (Millipore-Sigma), 10 μg/mL gentamicin (Gibco, Waltham, MA), and 0.05 mM β-mercaptoethanol (Millipore-Sigma). HEK293T cells were cultured in DMEM with L-glutamine and 4.5 g/L glucose (Millipore-Sigma) supplemented with 10% Serum Plus II Medium Supplement and 10 μg/mL gentamicin. To induce editing in BCBL-1 iCas9, cells were treated with 1 μg/mL doxycycline until the end of the growth curve. To induce lytic reactivation, cells were treated with 20 ng/mL TPA and 1 mM sodium butyrate for 2 d. To inhibit proteasome activity, cells were treated with 1.25 μM MG132 for 4 h.

### Primers and synthetic genes

All primers were purchased from Integrated DNA Technologies (Coralville, IA). Synthetic genes were purchased from Twist Biosciences (South San Francisco, CA). Sequences are found in **Table S2**.

### gRNAs

Primer sequences and cloning strategies for specific gRNA constructs are found in **Table S2**. gRNAs for the constitutive Cas9 backbone were cloned using annealed primers into BsmBI-linearized lentiGuide-Puro (a gift from Feng Zhang; Addgene catalog no. 52963). gRNAs for iCas9 were cloned into pLX-sgRNA (a gift from Eric Lander and David Sabatini; Addgene catalog no. 50662) as before [34]. To clone gRNAs targeting the BH3-only proteins, we modified the pRDA122_mKate backbone (a gift from the Genetic Perturbation Platform at the Broad Institute). First, the eGFP cDNA was PCR amplified from pLCE [53] using primers 229 and 230. Then, the eGFP PCR product or codon-optimized iRFP670 synthetic gene was cloned into a AgeI/BamHI-linearized pRDA122_mKate. The resulting gRNA vectors pRDA122_eGFP and pRDA122_iRFP670 now express eGFP and iRFP670, respectively, instead of mKate. The negative control gRNA SAFE4501 [40] was cloned into pRDA122_iRFP670. The negative control gAAVS1 and the rest of the targeting gRNAs were cloned into pRDA122_eGFP.

### cDNA cloning

All cDNAs used were cloned by Gibson assembly into pLC-coMCL1-IRES-hyg [24] using the NheI/BamHI restriction sites. pLC-mCherry-IRES-hyg is a gift from Eva Gottwein and was previously published [24]. coMARCHF5-3XFLAG cDNA was synthesized by Twist Biosciences. Site-directed mutagenesis were performed to construct the coMARCHF5-3XFLAG mutants using the pLC-coMARCHF5-3XFLAG WT-IRES-hyg as a template (**Table S2**).

### Lentiviruses and lentiviral infection

5 x 10^6^ HEK293T cells were plated in 10 cm dishes. The next day, the medium was replaced with DMEM without any supplements. 5 μg of transfer plasmid, 2.44 μg of pMD2.G (a gift from Didier Trono; Addgene catalog no. 12259) and 3.74 μg of pSPAX2 (a gift from Didier Trono; Addgene catalog no. 12260) were transfected with 39.13 μL of 1 mg/mL polyethylenimine (Sigma-Aldrich, 25kDa Mw branched, pH 7.4). After 4-6 h, media was replaced with complete RPMI-1640 with 10% Serum Plus II Medium Supplement.

Cells were grown in the cell culture incubator and viruses were harvested from the supernatant by filtering through a 0.45 μm polyethersulfone membrane.

Cells were either directly infected with undiluted lentiviral supernatants or spinfected with concentrated lentivirus at 1600 *g* for 45 min. After 24 h, cell culture media was replaced with fresh media with antibiotic selection (1 μg/mL puromycin or 100-250 μg/mL hygromycin).

### Growth curves

3 x 10^5^ cells were seeded in 1 mL in 12-well plates. Cells were counted by trypan blue exclusion using the Countess II Automated Cell Counter (Thermo Fisher) every 3 d. Cell densities were adjusted back to 3 x 10^5^ cells/mL to ensure that the cells did not undergo cell cycle arrest.

### Competition Assay

We first generated our single BH3-only protein KO in BC-3 Cas9 cells using a lentivirus expressing the targeting gRNA or a negative control gRNA SAFE4501. This lentivirus expresses the fluorescent reporter eGFP in its backbone. In parallel, we also engineered our control competitor BC-3 Cas9 cell line by transducing it with a second lentivirus expressing gAAVS1 and the fluorescent reporter iRFP670 instead of eGFP. After puromycin selection and confirmation of KO by Western blots, each eGFP+ KO cell line was mixed at 1:1 with iRFP670+ cells in 12-well plates. To establish an accurate baseline of eGFP:iRFP670 ratios, cell pools were analyzed by flow cytometry at this initial time point. Cells were then grown similar to our Growth Curve Assays and eGFP:iRFP670 ratios were continually monitored every 3 d for 15 d.

### Caspase 3/7 Assay

Twenty microliters of resuspended cells were dispensed into 96-well half-area microplates (Greiner Bio-One, Monroe, NC). An equal volume of reconstituted Caspase-Glo 3/7 Reagent (Promega, Madison, WI) was added to each sample and mixed by repeated pipetting. Samples were incubated in the dark for 30 min. Luminescence was read in a Synergy H1 microplate reader (Biotek, Winooski, VT) with an integration time of 2 s.

### Annexin V/7-AAD Staining

7 x 10^5^ cells were pelleted and washed twice in 0.5 mL 2% BSA in PBS. The cell pellets were resuspended in 100 μL Annexin V binding buffer (Biolegend, San Diego, CA) with 5 μL Annexin V-APC and/or 5 μL 7-AAD. Cells were stained in the dark for 15 min at 4°C. After incubation, an additional 350 μL of Annexin V binding buffer was added. Samples were analyzed on the BD LSR Fortessa.

### Western blots

Cells were lysed on ice for 10 min with radioimmunoprecipitation assay (RIPA) buffer (20 mM Tris-HCl at pH 7.5, 150 mM NaCl, 1 mM EDTA, 1 mM EGTA, 1% IGEPAL CA-630, 1% sodium deoxycholate) supplemented with protease inhibitor cocktail III (Millipore). Lysates were clarified by centrifuging at 12,000 g for 10 min at 4°C. Total protein concentrations from clarified lysates were determined using the Pierce BCA Protein Assay Kit (Thermo Scientific). Equal amounts of total protein were loaded and separated in bis-tris acrylamide gels in MES-SDS Running Buffer (Invitrogen) at 100 V. The proteins were transferred to a 0.2 µm nitrocellulose membrane in Towbin buffer (25 mM Tris, 192 mM glycine, 20% methanol, pH 8.3) for 1 h and 45 min at a constant current of 400 mA. Membranes were airdried, stained with Revert 700 Total Protein Stain (LICOR), and imaged using Odyssey Fc. Western blots were blocked with Intercept (TBS) blocking buffer (LICOR) diluted 1:2 in tris-buffered saline (TBS; 20 mM Tris, 150 mM NaCl, pH 7.6) for 1 h. The membranes were incubated with the primary antibody in TBS supplemented with 0.1% Tween-20 (TBST) overnight at 4°C (See **Table S2**).

The next day, the membranes are washed 5x with TBST for 7 min each before incubating with IR800CW secondary antibody (LICOR) for 1 h. Final washes in TBST were done before imaging on the Odyssey M (LICOR).

### Reciprocal Co-Immunoprecipitation (co-IP)

For each co-IP reaction, 2 x 10^7^ BC-3 cells were washed thrice with phosphate-buffered saline (PBS, 137 mM NaCl, 2.7 mM KCl, 1.47 mM KH_2_PO_4_, 8.1 mM Na_2_HPO_4_, pH 7.4) then lysed with 600 μL IP buffer (25 mM HEPES, 100 mM NaCl, 10 mM CaCl_2_, 5 mM MgCl_2_) freshly supplemented with 1% digitonin and protease inhibitor cocktail III. To 400 μL of clarified lysate, IP antibodies were added and incubated on a rotisserie overnight at 4°C: 3.5 μL anti-NOXA antibody (0.55 mg/mL), 4.5 μL anti-MARCHF5 antibody (0.21 mg/mL) or 0.77 μL rabbit IgG (2.5 mg/mL). The remaining lysates were stored at -80°C as input. The following day, 8 μL of Protein A/G magnetic beads (Pierce) were added to each IP and reincubated on a rotisserie for 4.5 h at 4°C. The beads were washed twice with 500 μL IP buffer with 0.1% digitonin and resuspended in 45 μL 1.5X LDS loading buffer (40 mM Tris-Cl, 53 mM Tris Base, 0.75% lithium dodecyl sulfate, 3.75% glycerol, 0.2 mM EDTA, 3.75% β- mercaptoethanol, pH 8.5). For controls, 27 μL of input was mixed with 9 μL 4X LDS loading dye. All samples were denatured at 90°C for 7 min. 36 μL of input and 20 μL of each IP were run in each lane of a 12% bis-tris acrylamide gel in MES-SDS Running Buffer at 80V. Western blots for MARCHF5 or NOXA were performed as above.

## ACKNOWLEDGMENTS

This work is supported by a Transition Career Development Award K22 CA241355 from the National Cancer Institute of the National Institutes of Health (NIH) to M.M. Additional support was provided by the Center for Microbial Pathogenesis and Host Inflammatory Responses program grant P30 GM103625 from the National Institute of General Medical Sciences (NIGMS) of the NIH and the Winthrop P. Rockefeller Cancer Institute at UAMS. J.G., A.N., and A.F. were supported by the Undergraduate Summer Research Fellowship from the Arkansas IDeA Network of Biomedical Research Excellence (INBRE) funded by P20 GM103429 from NIH/NIGMS. We thank Todd Spears for initial co-immunoprecipitation experiments. The authors declare no conflicts of interest. The funders had no role in study design, data collection and analysis, decision to publish, or preparation of the manuscript.

## AUTHOR CONTRIBUTIONS

Conceptualization, Funding Acquisition, Supervision, Writing – original draft: M.M., Writing – review and edits: M.M., J.R.B., P.V., K.W.B. Investigation, Formal Analysis, Methodology, Validation: M.M., P.V., J.R.B., J.G., A.N., A.F., D.D. Resources: K.W.B.

